# Selective loss of Primary Cilia and Neurotrophic Signaling in G51D α-Synuclein Mice Highlights a Common Pathway to Parkinson’s Disease

**DOI:** 10.64898/2026.02.25.708014

**Authors:** Yu-En Lin, Ebsy Jaimon, YoungDoo Kim, Annabeth Loftman, Aaran Vijayakumaran, Benjamin D.W. Belfort, Claire Y. Chiang, Benjamin R. Arenkiel, Huda Y. Zoghbi, Suzanne R. Pfeffer

## Abstract

Parkinson’s disease is characterized by dopaminergic neuron loss and accumulation of α-synuclein aggregates in the brain. G51D α-synuclein knock-in mice provide a genetically and clinically relevant model of disease, exhibiting early olfactory deficits, age-dependent motor impairment, and progressive phospho-α-synuclein accumulation. In multiple Parkinson’s disease models, striatal cholinergic and parvalbumin interneurons, as well as astrocytes, lose primary cilia and the neurotrophic signaling needed to sustain dopaminergic neurons. We show here that G51D-α-synuclein mice share these phenotypes. Phospho-Ser129 α-synuclein accumulation correlates with cilia loss in cholinergic interneurons but not in medium spiny neurons that accumulate higher phospho-α-synuclein levels. In the piriform cortex, parvalbumin neurons lose primary cilia and downregulate Neurturin, potentially contributing to olfactory dysfunction. Within the peripheral olfactory epithelium, horizontal basal cells lose cilia, whereas multi-ciliated olfactory sensory neuron cilia remain intact. These findings reveal convergent cellular vulnerabilities across Parkinson’s disease models and highlight a pathogenic role for impaired ciliary signaling.

**Teaser:** Loss of primary cilia may contribute to dopamine neuron loss in both inherited and common Parkinson’s disease.

## Introduction

Most Parkinson’s disease (PD) is of unknown cause, although pesticides and other environmental insults are strongly implicated (*1*). Hallmarks of PD include resting tremor, rigidity, and bradykinesia due to death of dopaminergic neurons in the substantia nigra that govern movement coordination (*1*). Loss of smell and constipation can precede these symptoms by more than 10 years, along with rapid eye movement (REM) sleep behavior disorder in which a person acts out their dreams because normal, REM-related muscle paralysis is absent. Abnormal aggregation of alpha-synuclein (α-Syn) is a key contributor to PD, promoting neurodegeneration through toxic oligomers and fibrils that disrupt synaptic function, membrane integrity, and cellular homeostasis (*2, 3*). Misfolded α-Syn spreads between neurons, leading to so-called Lewy pathology (*4, 5*).

10-15% of PD cases are caused by identifiable genetic mutations or genetic risk variants, including missense mutations in *SNCA* encoding α-Syn, activating mutations in the Leucine Rich Repeat Kinase 2 (LRRK2) or loss of function mutations in *GBA1* encoding lysosomal glucocerebrosidase (*6*). In our efforts to understand the molecular basis of LRRK2-PD, we showed that LRRK2 phosphorylation of Rab10 protein blocks primary cilia formation in cell culture by a process that requires the phosphoRab effector protein, RILPL1 (*7, 8*). In the striatum of mice harboring pathogenic LRRK2 mutations (G2019S, R1441C) or lacking the genes encoding either PPM1H or PPM1M phosphatases that reverse LRRK2 action (*9, 10*), we have shown that primary cilia are lost in a cell-type specific manner: rare, cholinergic (ChAT) and parvalbumin (PV) neurons and ALDH1L1^+^ or GFAP^+^ astrocytes lose their cilia, while the much more abundant medium spiny neurons do not (*8, 11–15*). Cell type specificity of cilia loss is not dictated by overall LRRK2 or PPM1H expression levels (*12*). However, among ChAT neurons, higher LRRK2 levels correlate with decreased ciliation (*12*).

Primary cilia are essential for Hedgehog signaling (*16*), which is critical for the survival of nigral dopamine neurons that project to the dorsal striatum and the cholinergic neurons located there (*17*). We have shown that cilia loss correlates with loss of Hedgehog signaling in the striatum (*8*), and the concomitant loss of cilia-dependent transcription of neurotrophic factors that support dopamine neurons (*12, 13, 15*). Thus, in the mouse striatum, cilia loss decreases production of glial derived neurotrophic factor (GDNF) by ChAT neurons (*12*), Neurturin (NRTN) by PV neurons (*13*), and Brain derived neurotrophic factor (BDNF) by astrocytes (*18*). Loss of these factors leads to decreased density of the dopaminergic axonal arbor in the striatum, as monitored by loss of tyrosine hydroxylase or GDNF Receptor Alpha-1 staining (*15*). The importance of Hedgehog signaling is recapitulated in *Gba1* mutant mice that retain their cilia but show decreased Hedgehog signaling because of an altered lipid composition in their otherwise normal-appearing cilia (*18*).

Unlike most idiopathic PD, ∼30% of LRRK2-PD lack classical Lewy body pathology (*19*). Because of this, it was not clear whether LRRK2-PD pathogenesis was similar to that of idiopathic disease. Very recently, Jensen and colleagues (*20*) identified a non-inclusion, aggregated α-Syn in previously considered, “non-Lewy body” LRRK2 cases. These results suggest that Lewy-body negative LRRK2-PD is not associated with a lack of α-Syn aggregation in neurons, but rather a deficiency in the appearance of inclusions.

In our prior analyses of postmortem human striatum from patients with LRRK2 mutations, idiopathic PD, or age matched controls, we detected the same cell type-specific, striatal primary cilia loss in patients with LRRK2 pathway mutations, and also in patients with idiopathic disease (*12, 13*). This suggested that synuclein-driven pathology might also trigger cell-type specific cilia loss. To explore this further, we sought a physiologically relevant mouse model for synuclein disease. A recently generated *Snca^G51D/G51D^* (G51D) mouse, where the mutation is knocked into the endogenous murine *Snca* locus (*21*) provided an ideal model. These mice express mutant α-Syn^G51D^ in the native spatiotemporal pattern and display the cascade of molecular events and symptom progression characteristic of human PD, including early loss of olfaction and enteric dysfunction followed by age-related motor symptoms (*21*). We show here that like LRRK2 pathway mutant animals, G51D mice show cell-type selective cilia loss and loss of striatal neuroprotection.

## Results

### Loss of cilia in vulnerable striatal interneurons and astrocytes of G51D mice

We first examined the ciliation status of neurons in the striatum of WT and G51D mice. Figure 1 shows examples of ChAT neurons (Fig. 1, A-D), PV neurons (Fig. 1, E-H) and ALDH1L1^+^ astrocytes (Fig. 1, I-L) and their primary ciliation status, in the dorsolateral striatum at 3 and 12 months of age (3M, 12M). Neuronal cilia are labeled with anti-adenylate cyclase III while astrocyte cilia are visualized using anti-Arl13b antibodies. As shown in the quantitation at right, when compared with age-matched, wild type mice, all cell types showed cilia loss even at 3 months of age, although the astrocytes appeared to require aging before a highly significant difference was detected. Nevertheless, all cell types analyzed showed a greater loss of primary cilia with age in this G51D mouse model.

**Fig. 1.**
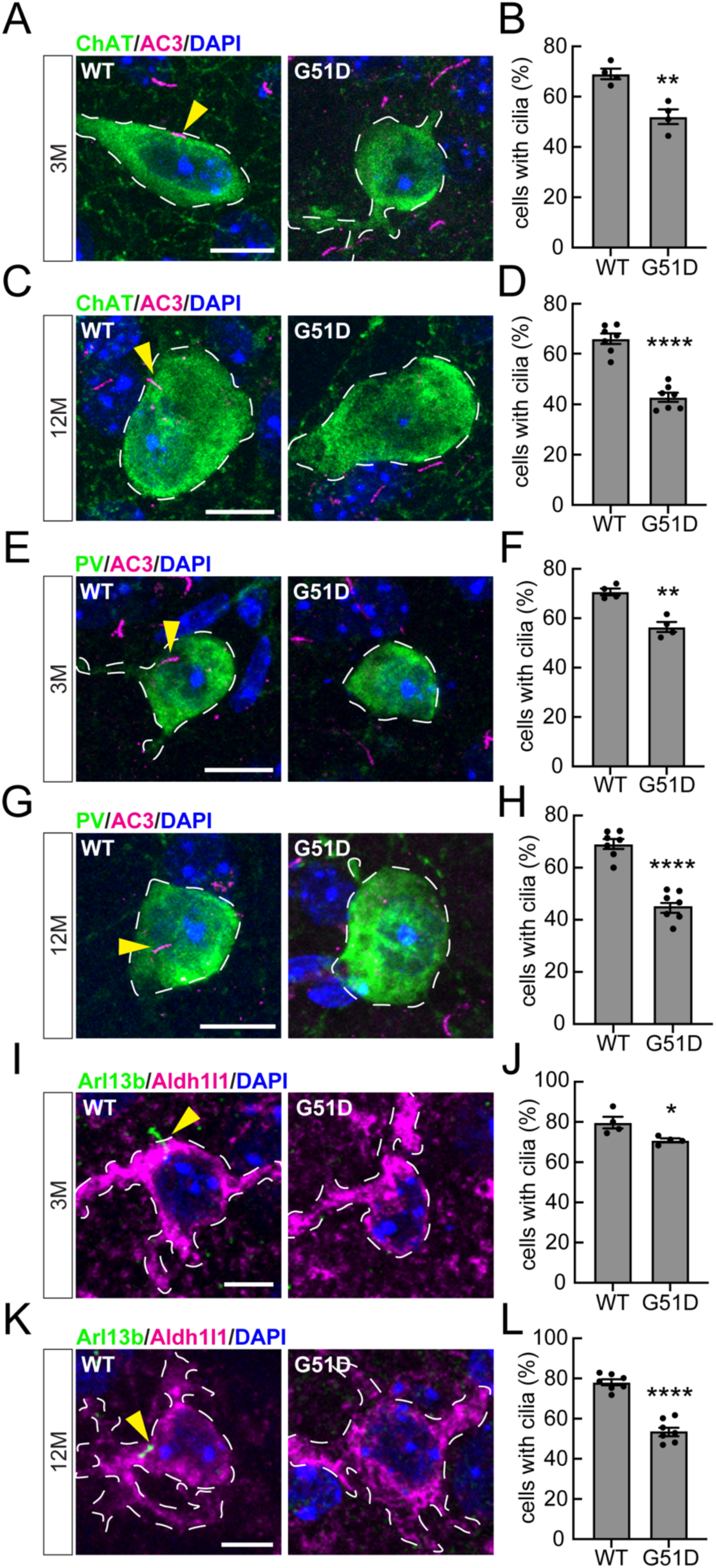
The *Snca^G51D/G51D^* mutation induces age-dependent cilia defects in vulnerable cells of the mouse dorsal striatum. **(A, C, E, G, I, K)** Representative confocal immunofluorescence images of dorsal striatal sections from 3- and 12-month-old (3M and 12M) wild type and *Snca^G51D/G51D^* mice. Cholinergic and parvalbumin (PV) interneurons were labeled with anti-ChAT and anti-PV antibodies (green), respectively. Primary cilia were labeled with anti-AC3 antibody (magenta; yellow arrowheads). Nuclei were labeled with DAPI (blue). Astrocytes were labeled with anti-Aldh1l1 antibody (magenta), and astrocytic primary cilia were labeled with anti-Arl13B antibody (green; yellow arrowheads). Scale bars, 10 μm for cholinergic and PV neurons; 5 μm for astrocytes. **(B and D)** Quantitation of ChAT^+^ neurons containing a cilium in 3- and 12-month-old wild type and *Snca^G51D/G51D^* mice, respectively. **(F and H)** Quantitation of PV neurons containing a cilium in 3- and 12-month-old wild type and *Snca^G51D/G51D^*mice, respectively. **(J and L)** Quantitation of Aldh1l1^+^ astrocytes containing a cilium in 3- and 12-month-old wild type and *Snca^G51D/G51D^* mice, respectively. Values represent mean ± SEM from four (3-month-old) or seven (12-month-old) mice per group, with two to three sections analyzed per mouse. More than 37 ChAT^+^ neurons, 40 PV neurons and 56 Aldh1l1^+^ astrocytes were scored per mouse. Statistical significance was determined using an unpaired Student’s *t* test. **p = 0.0033 for ChAT^+^ neurons at 3 months; ****p < 0.0001 for ChAT^+^ neurons at 12 months; **p = 0.0012 for PV neurons at 3 months; ****p < 0.0001 for PV neurons at 12 months; *p = 0.0291 for Aldh1l1^+^ astrocytes at 3 months; ****p < 0.0001 for Aldh1l1^+^ astrocytes at 12 months.

Similar to what we have previously reported for mouse genetic models and for patients with idiopathic or genetic PD, medium spiny neurons identified using anti-DARPP32 antibodies showed normal ciliation, even after 12 months, despite expression of G51D α-Syn (fig. S1). These data confirm the relative resilience of the cilia of medium spiny neurons to this mutant synuclein insult.

### Loss of striatal neuroprotective factors

Loss of cilia would be predicted to lead to loss of transcription of cilia- and Hedgehog-dependent neurotrophic factors (*16*). Figures 2-4 show RNAscope fluorescence in situ hybridization (FISH) to monitor expression of the neurotrophic factors, GDNF (Fig. 2), NRTN (Fig. 3) and BDNF (Fig. 4) in specific cell types in the dorsolateral striatum. Figure 2 shows examples of ciliated and non-ciliated ChAT neurons in wild type and G51D mice; control labeling in the absence of the specific FISH probe is shown at top right (Fig. 2A). Quantitation of these data revealed that G51D mice express fewer GDNF transcripts than wild type mice, with the decrease proportional to their cilia loss at 12 months of age, ∼40% (Figs. 2B, 1D). Moreover, loss of GDNF expression was cilia-dependent (Fig. 2C). Figure 3 shows similar analysis of PV neurons (Fig. 3, A and B); again, cilia loss led to significant loss of NRTN transcripts in striatal PV neurons (Fig. 3B) and expression was cilia dependent (Fig. 3C). Finally, analysis of ALDH1L1^+^ astrocytes in this brain region showed strong loss of BDNF production (Fig. 4, A and B); in this case, cilia dependence was less clear (Fig. 4C), although we have shown BDNF to be a cilia-dependent gene in prior studies of 5-month-old mice (*18*).

**Fig. 2.**
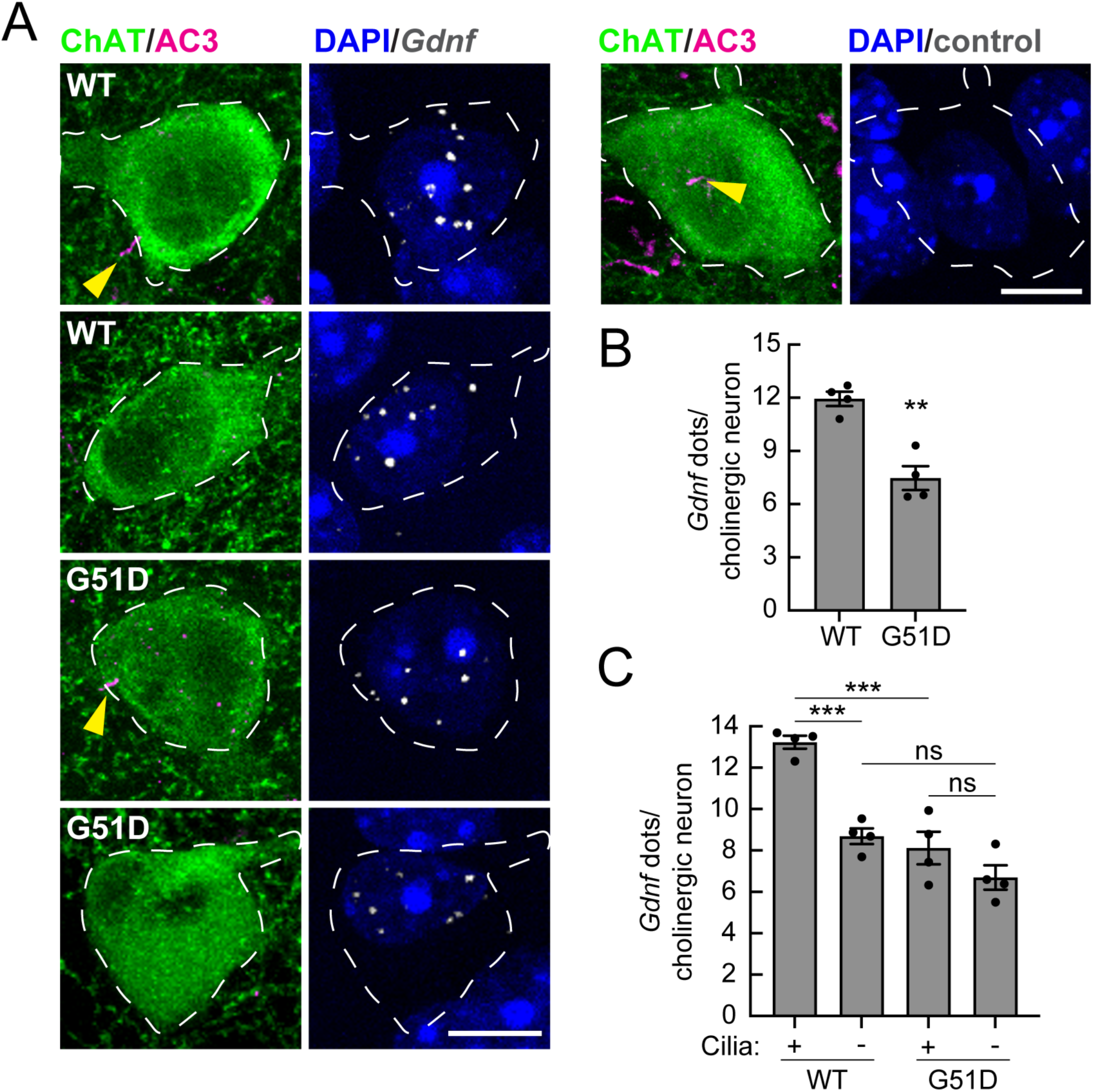
Reduced cilium-dependent GDNF expression in striatal cholinergic neurons of Snca*^G51D/G51D^* mice. **(A)** Representative confocal immunofluorescence images of cholinergic neurons from 12-month-old wild type and *Snca^G51D/G51D^* mice. Cholinergic interneurons were labeled with anti-ChAT antibody (green), primary cilia were labeled with anti-AC3 antibody (magenta; yellow arrowheads), and nuclei were labeled with DAPI (blue). Sections were subjected to RNAscope in situ hybridization using a *Gdnf* RNA probe (white dots). Images of a control neuron processed without the RNAscope probe are shown above panel (B), at the top right. Scale bar, 10 μm. **(B)** Quantitation of the average number of *Gdnf* dots per ChAT^+^ cholinergic interneuron. **(C)** Quantitation of the average number of *Gdnf* dots segregated by ciliation status. Values represent mean ± SEM from four mice per group, with two to three sections analyzed per mouse. More than 49 ChAT^+^ neurons were scored per mouse. Statistical significance was determined using an unpaired Student’s *t* test or one-way ANOVA with Tukey’s post hoc test. **p = 0.0013 (wild type versus G51D); ***p = 0.0004 (ciliated wild type versus unciliated wild type); ***p = 0.0001 (ciliated wild type versus ciliated G51D). Differences were not significant (ns; p = 0.0983 for unciliated wild type versus unciliated G51D; p = 0.3053 for ciliated G51D versus unciliated G51D).

**Fig. 3.**
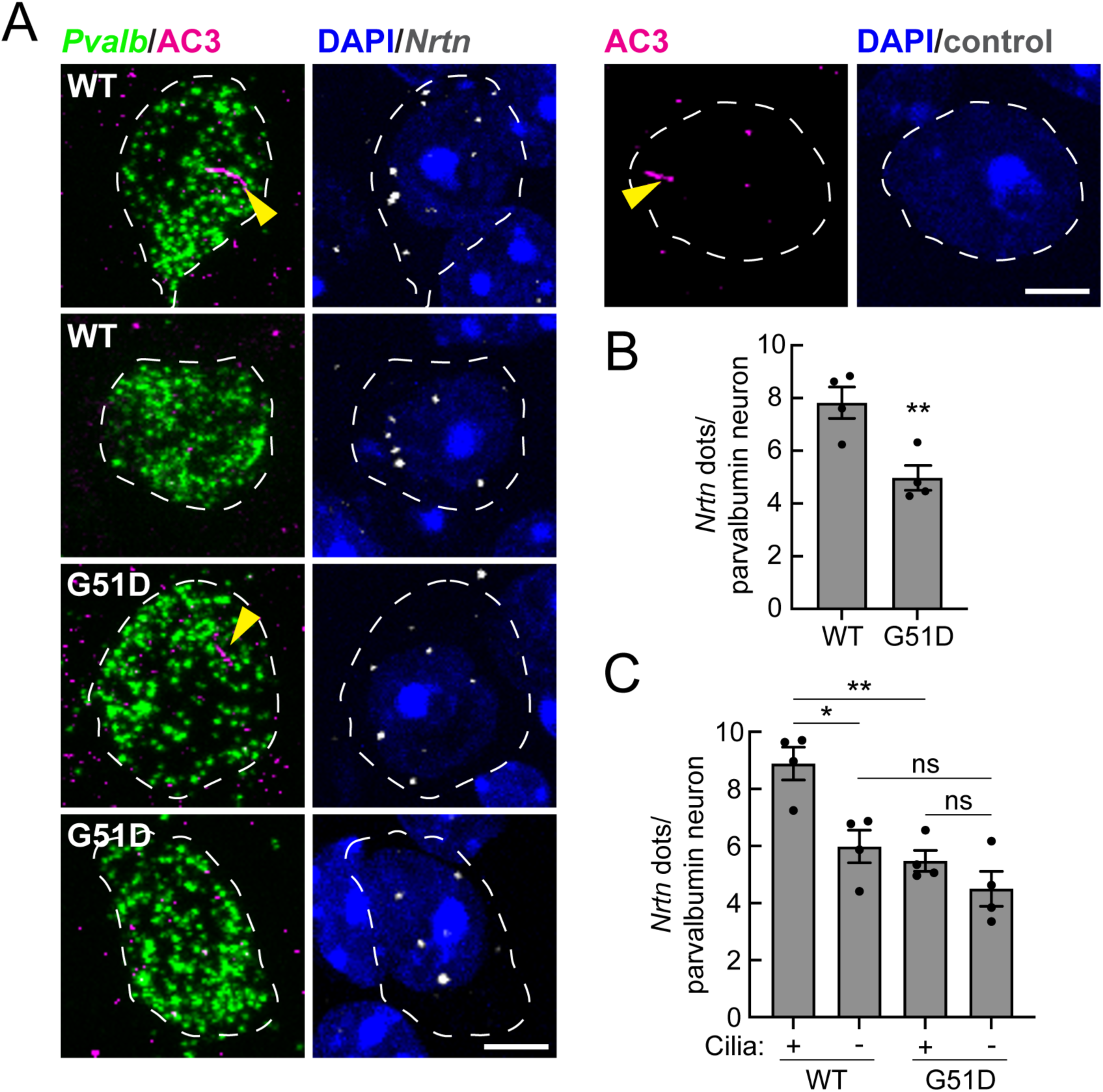
Reduced cilium-dependent Neurturin expression in striatal parvalbumin neurons of Snca*^G51D/G51D^* mice. **(A)** Representative confocal immunofluorescence images of parvalbumin (PV) neurons from 12-month-old wild type and *Snca^G51D/G51D^* mice. PV interneurons were identified using an RNAscope probe for *parvalbumin* (green). Primary cilia were labeled with anti-AC3 antibody (magenta; yellow arrowheads), and nuclei were labeled with DAPI (blue). Sections were subjected to RNAscope in situ hybridization using a *Nrtn* RNA probe (white dots). Images of a control neuron processed without the RNAscope probe are shown above panel (B), at the top right. Scale bar, 5 μm. **(B)** Quantitation of the average number of *Nrtn* dots per PV interneuron. **(C)** Quantitation of the average number of *Nrtn* dots segregated by ciliation status. Values represent mean ± SEM from four mice per group, with two to three sections analyzed per mouse. More than 49 PV neurons were scored per mouse. Statistical significance was determined using an unpaired Student’s *t* test or one-way ANOVA with Tukey’s post hoc test. **p = 0.0091 (wild type versus G51D); *p = 0.0115 (ciliated wild type versus unciliated wild type); **p = 0.0037 (ciliated wild type versus ciliated G51D). Differences were not significant (ns; p = 0.2625 for unciliated wild type versus unciliated G51D; p = 0.5898 for ciliated G51D versus unciliated G51D).

**Fig. 4.**
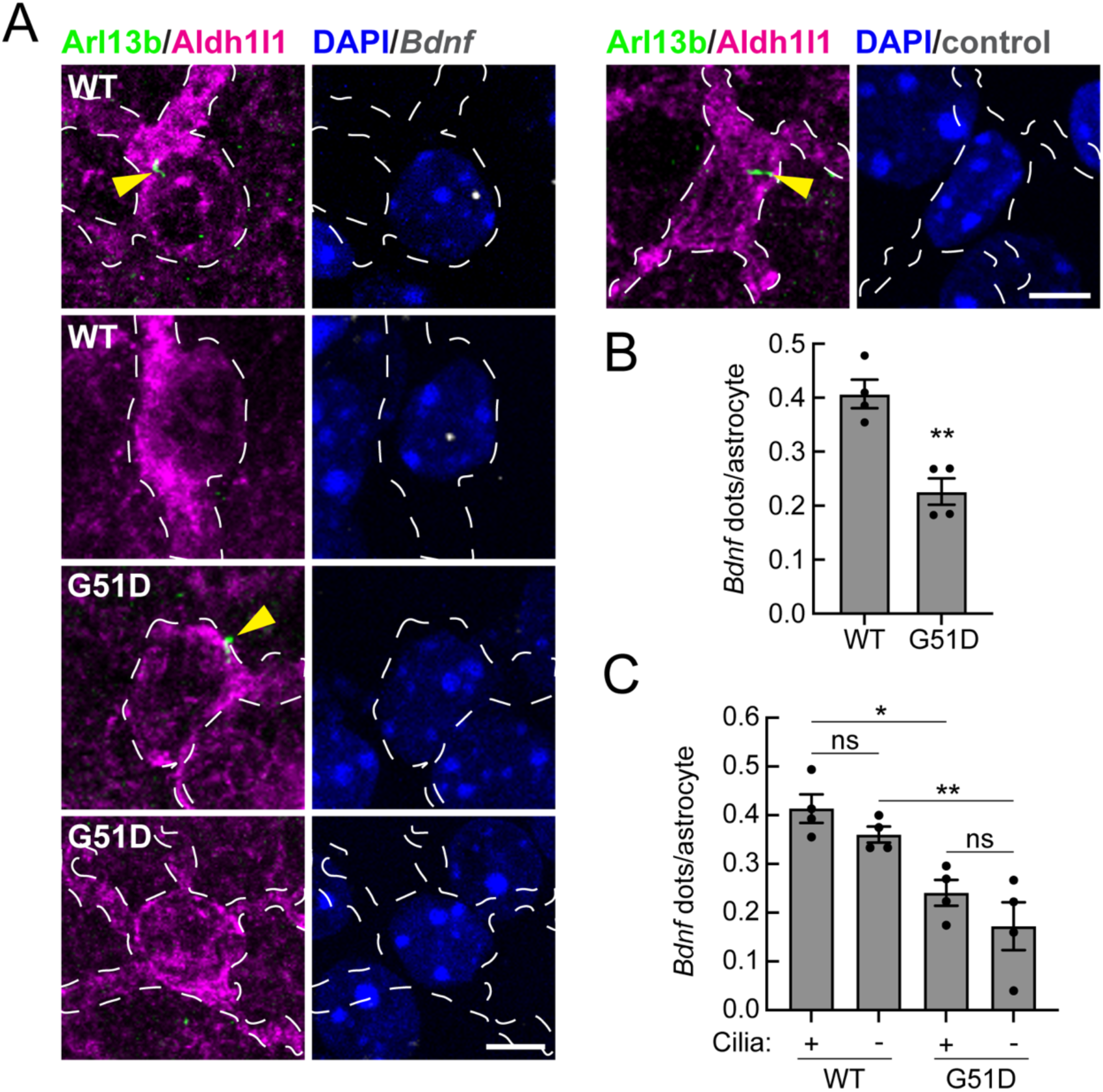
Reduced BDNF expression in striatal astrocytes of *Snca^G51D/G51D^* mice. **(A)** Representative confocal immunofluorescence images of astrocytes from 12-month-old wild type and *Snca^G51D/G51D^* mice. Astrocytes were labeled with anti-Aldh1l1 antibody (magenta), and astrocytic primary cilia were labeled with anti-Arl13B antibody (green; yellow arrowheads), and nuclei were labeled with DAPI (blue). Sections were subjected to RNAscope in situ hybridization using a *Bdnf* RNA probe (white dots). Images of a control astrocyte processed without the RNAscope probe are shown above panel (B), at the top right. Scale bar, 5 μm. **(B)** Quantitation of the average number of *Bdnf* dots per Aldh1l1^+^ astrocyte. **(C)** Quantitation of the average number of *Bdnf* dots segregated by ciliation status. Values represent mean ± SEM from four mice per group, with two to three sections analyzed per mouse. More than 79 Aldh1l1^+^ astrocytes were scored per mouse. Statistical significance was determined using an unpaired Student’s *t* test or one-way ANOVA with Tukey’s post hoc test. **p = 0.0024 (wild type versus G51D); *p = 0.0128 (ciliated wild type versus ciliated G51D); **p = 0.0072 (unciliated wild type versus unciliated G51D). Differences were not significant (ns; p = 0.6649 for ciliated wild type versus unciliated wild type; p = 0.4751 for ciliated G51D versus unciliated G51D).

### Loss of primary cilia in the Piriform cortex

An important prodromal symptom of PD is loss of olfaction (*22*). G51D mice show phosphoSer129 (pS129) α-Syn accumulation in the piriform cortex that likely contributes to their olfactory deficits (*21*). PV interneurons in this brain region are thought to be important inhibitory gatekeepers that shape how odor information is processed and learned (*23–25*). In the piriform cortex of wild type, 3-month-old mice, PV neurons were 50% ciliated (Fig. 5, A and B) compared with 70% ciliated in the dorsolateral striatum at this age (Fig. 1F). At 12 months, these values decreased to 40% ciliated in the wild type piriform cortex (Fig. 5, C and D) compared with 70% in the striatum (Fig. 1H). Thus, even in wild type mice, cilia are more likely to be lost with age in this brain region.

**Fig. 5.**
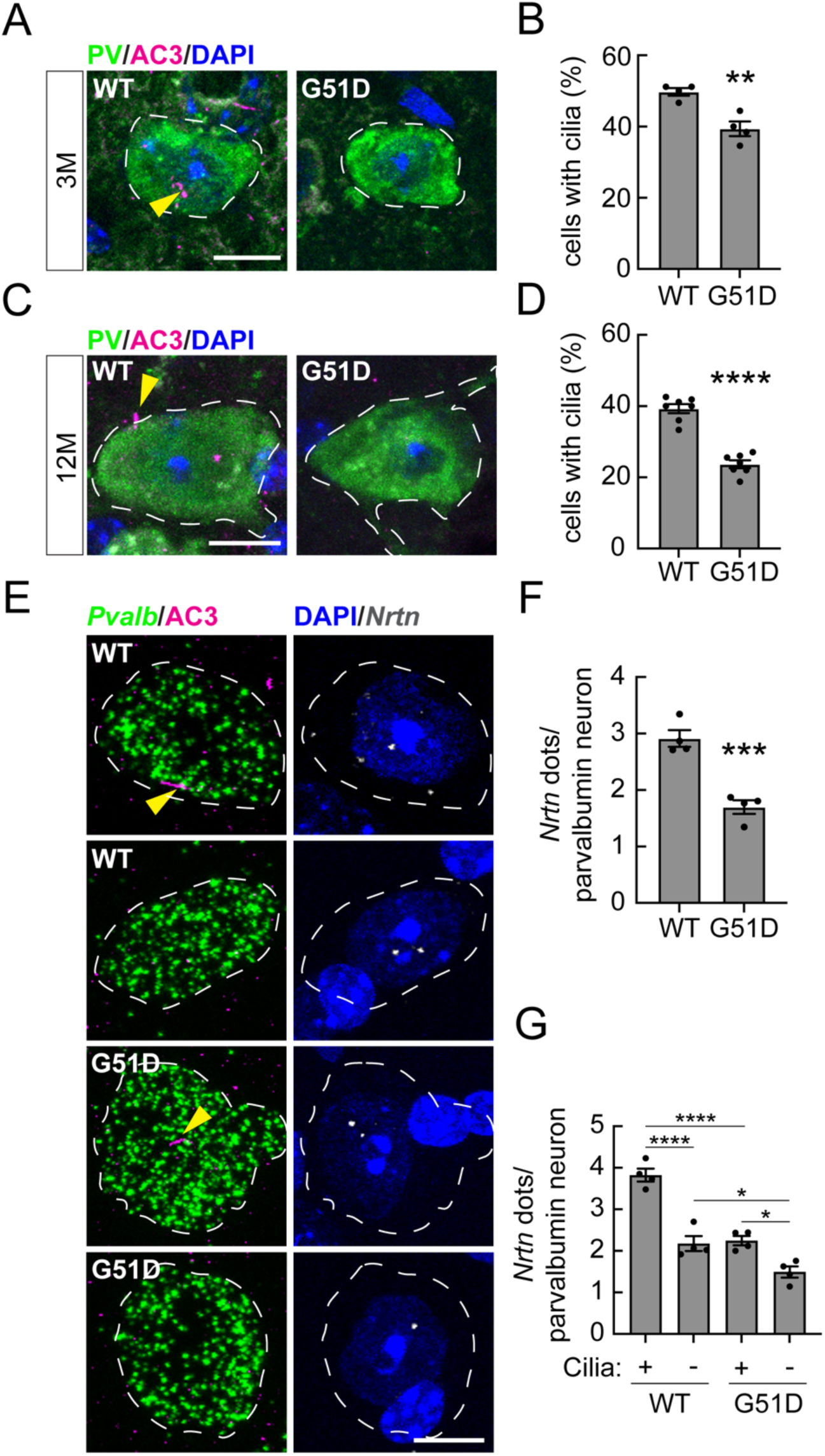
The *Snca^G51D/G51D^*mutation induces age-dependent cilia defects and reduces Neurturin expression in parvalbumin neurons of the piriform cortex. **(A and C)** Representative confocal immunofluorescence images of parvalbumin (PV) neurons from 3- and 12-month-old wild type and *Snca^G51D/G51D^* mice. PV interneurons were labeled with anti-PV antibody (green), primary cilia were labeled with anti-AC3 antibody (magenta; yellow arrowheads), and nuclei were labeled with DAPI (blue). Scale bars, 10 μm. **(B and D)** Quantitation of PV neurons containing a cilium in 3- and 12-month-old wild type and *Snca^G51D/G51D^* mice, respectively. **(E)** Representative confocal images of PV neurons from 12-month-old wild type and *Snca^G51D/G51D^*mice. Primary cilia were labeled with anti-AC3 antibody (magenta; yellow arrowheads), and nuclei were labeled with DAPI (blue). Sections were subjected to RNAscope in situ hybridization using a *Nrtn* and *Pvalb* RNA probe (white or green dots). Scale bars, 10 μm. **(F)** Quantitation of the average number of *Nrtn* dots per PV interneuron. **(G)** Quantitation of the average number of *Nrtn* dots segregated by ciliation status. Values represent mean ± SEM from four mice per group, except for Fig. 5D, which includes seven (12-month-old) mice for the ciliation analysis, with two to three sections analyzed per mouse. More than 43 PV neurons were scored per mouse. Statistical significance was determined using an unpaired Student’s *t* test or one-way ANOVA with Tukey’s post hoc test. **p = 0.0040 for PV neurons at 3 months; ****p < 0.0001 for PV neurons at 12 months; ***p = 0.0007 (wild type versus G51D); ****p < 0.0001 (ciliated wild type versus unciliated wild type; ciliated wild type versus ciliated G51D); *p = 0.0285 (unciliated wild type versus unciliated G51D); *p = 0.0155 (ciliated G51D versus unciliated G51D).

In G51D mice, as early as 3 months, but more significantly at 12 months, PV neurons lost almost half of their primary cilia, on top of the loss due to aging (Fig. 5, A-D). At 6 months, G51D mice show olfactory deficits (*21*), so one might have predicted that cilia loss would have been greater in PV neurons in this brain region compared with the striatum. NRTN expression in PV neurons is significantly decreased by 12 months, in direct proportion to ciliary loss (Fig. 5, E and F). Again, NRTN production was cilia-dependent, similar to that in the striatum (Fig. 3C), and as we have previously reported in other genetic backgrounds (Fig. 5G; (*13, 15*)). Absolute NRTN transcript levels were lower in wild type piriform cortex compared with wild type striatum, measured in the same tissue sections (∼8 versus ∼3 dots per cell). Beyond general neuroprotection, the full consequences of PV neuron cilia loss in both piriform cortex and striatum remain to be established.

### Analysis of cilia loss in the olfactory epithelium

Peripheral olfactory processing begins in the olfactory epithelium, where olfactory sensory neurons (OSNs) detect odorants and project their axons to the olfactory bulb (*26, 27*). OSNs are multi-ciliated neurons in which olfactory G protein–coupled odorant receptors are localized to the sensory cilia (*28, 29*). Each OSN expresses just one out of ∼400 different odorant receptors, and all sensory neurons expressing a given receptor project to a unique pair of glomeruli in the olfactory bulb (*29, 30*). Other ciliated cells in the olfactory epithelium are the horizontally oriented basal cells, which are singly ciliated, quiescent reserve stem cells that become activated after injury to regenerate both OSNs and non-neuronal epithelial cell types (*31*). Joiner et al. (*32*) demonstrated that horizontal basal cells require their primary cilia to support regeneration of the olfactory epithelium following injury.

To gain further insight into the loss of olfaction in G51D mice, we also explored the ciliation status of their olfactory epithelial cells. Figure 6A shows staining of horizontal basal cells, labeled with anti-p63 and their single, primary cilia (labeled with anti-Arl13B), in wild type and G51D α-Syn olfactory epithelium. Quantification of ciliation status and cell number showed ∼30% loss of cilia (Fig. 6B) with no significant loss of cell number (Fig. 6C). Cilia loss will decrease the ability of these cells to respond to Hedgehog and other growth and differentiation signals needed during regeneration processes (*32*).

**Fig. 6.**
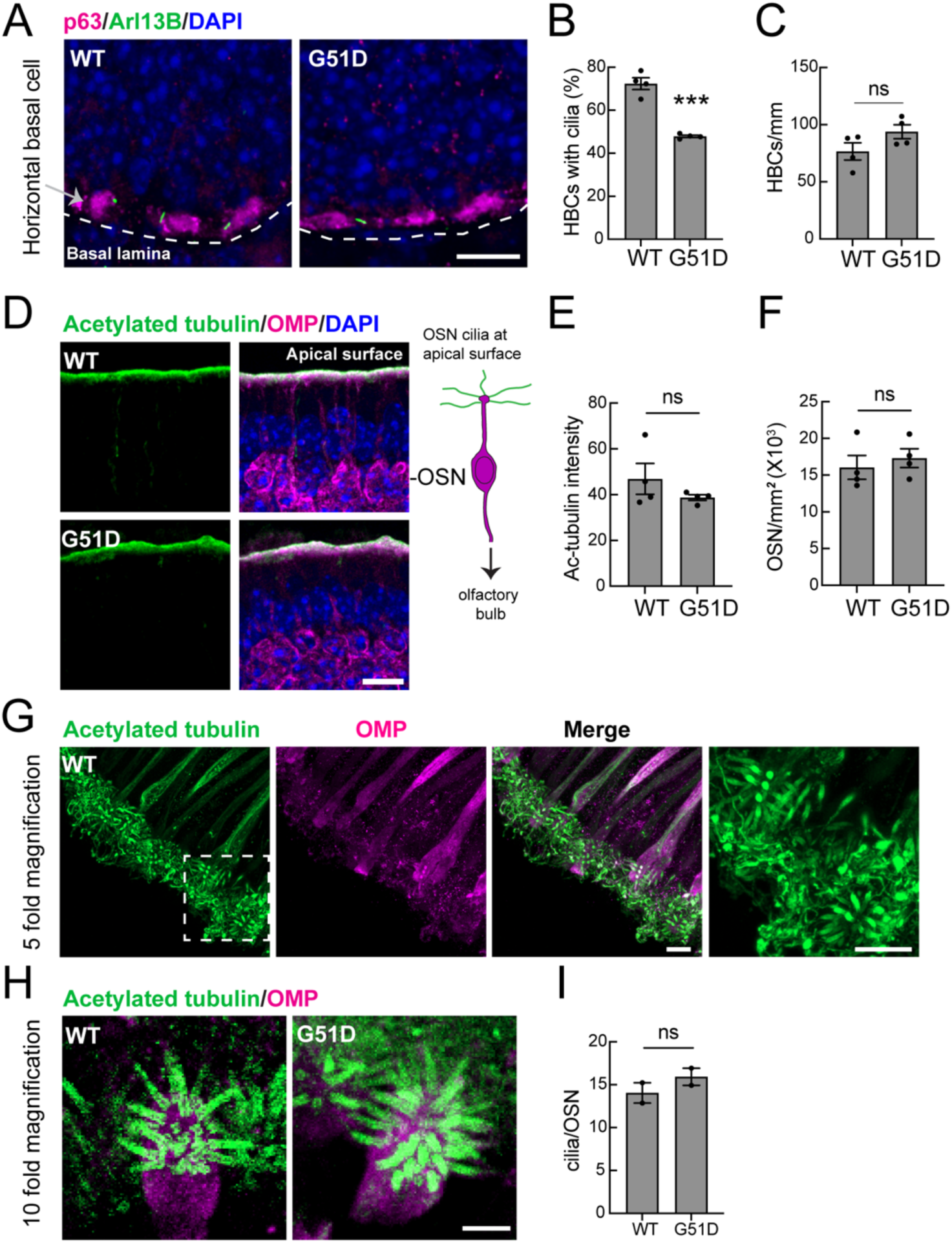
Primary cilia loss in the olfactory stem cells of 12-month Snca*^G51D/G51D^* mice. Representative images of horizontal basal cells (HBCs) from the olfactory epithelium of 12-month-old wild type and Snca*^G51D/G51D^* mice. HBCs were labeled with anti-p63 antibody (magenta), primary cilia with anti-Arl13B antibody (green), and nuclei with DAPI (blue). Dashed lines indicate the basement membrane. Scalebar, 10 µm. **(B)** Quantification of the percentage of HBCs containing a cilium. >100 HBCs were scored per mouse from 2-3 sections. **(C)** Quantification of HBCs per mm of the olfactory epithelium. >25 fields were analyzed per animal. **(D)** Representative images of the olfactory epithelium of 12-month-wild type and *Snca^G51D/G51D^* mice. Olfactory sensory neurons (OSNs) in D, G, and H were labeled with anti-OMP antibody (magenta), cilia with anti-Acetylated alpha-tubulin (green), and nuclei with DAPI (blue). Scale bar, 10 µm. **(E)** Quantitation of intensity of acetylated tubulin at the apical surface of the olfactory epithelium. >25 fields were analyzed per animal. **(F)** Quantitation of OSNs. > 15 fields were analyzed per animal. **(G)** Representative confocal images of the OSN cilia of 12-month wild type mice visualized using expansion microscopy. Scale bar, 2 µm. **(H)** Representative confocal images of the OSN cilia of 12-month-old wild type and *Snca^G51D/G51D^* mice visualized using expansion microscopy. Scale bar, 1 µm. **(I)** Quantitation of cilia per OSN. >15 OSNs were scored per animal. Values represent mean ± SEM from four mice per group for Fig. 6B, C, E, and F, and from two mice per group for Fig.6I. Statistical significance was determined using an unpaired Student’s *t* test. ***p = 0.0001 for WT versus *Snca^G51D/G51D^*mice.

OSNs were identified using anti-olfactory marker protein (OMP) antibodies; apically localized cilia were labeled with anti-acetylated tubulin (Fig. 6D). Ciliary staining was restricted to the apical surface of the olfactory epithelium, but individual cilia could not be resolved by conventional fluorescence microscopy (Fig. 6D). Nevertheless, the intensity of staining was not different in these micrographs between wild type and G51D mice (Fig. 6E) and the overall number of OSNs was also unchanged (Fig. 6F). To increase the resolution of our analyses, we incorporated 4X expansion microscopy and Airyscan super-resolution to enable visualization of entire, multi-ciliated OSN tufts (Fig. 6, G and H; fig. S2). Although it was sometimes complicated to count, precisely, the number of individual cilia within each multi-ciliated tuft, we did not discern any obvious differences between the multi-ciliation status of OSNs, when comparing wild type and G51D α-Syn olfactory epithelium (Fig. 6I). Note that the molecular mechanisms that generate multiple cilia on a single OSN differ from those that underlie formation of the more common single primary cilium (*33–35*), so it is perhaps not unexpected that these cell types respond differently to the G51D mutation. Altogether, these data suggest that olfactory regeneration may be impacted by loss of cilia on horizontal basal cells; further experiments will be needed to pinpoint more precisely, the full defects underlying olfactory dysfunction in this synucleinopathy model.

### Overall LRRK2 pathway activation appears unchanged

Because LRRK2 activation and Rab GTPase phosphorylation are known to block primary cilia formation (*8, 36, 37*), one possible explanation for cilia loss in the G51D mouse model is that α-Syn aggregates cause lysosome stress, thereby activating LRRK2 kinase (*38–40*). To test this, we harvested brains from 9-month-old mice and analyzed the tissue by western blot (fig. S3). We failed to detect significant differences in levels of pRab10, pRab12, or LRRK2 kinase when comparing wild type and G51D mice at the level of whole brain tissue. It is possible that LRRK2 is only activated in select cell types that are vulnerable to cilia loss, in which case such activation would not be able to be detected in a whole brain lysate. Indeed, even in a mouse where LRRK2 is inappropriately activated in all cell types, increases in pRab12 are ∼25% in overall magnitude in whole brain tissue (cf. (*14*)).

### PhosphoSer129 synuclein levels correlate with cilia loss in vulnerable cells

Phosphorylation of α-synuclein at Serine 129 (pS129-α-Syn) is a major posttranslational modification seen in postmortem human PD tissue (*41*). In G51D mice, pS129-α-Syn is detected in the motor cortex and structures necessary for motor coordination, including the striatum, pons, brainstem, and spinal cord (*21*). We evaluated pS129-α-Syn levels in the striatum, comparing ChAT neurons that are vulnerable to cilia loss with medium spiny neurons that are not. G51D mice showed much higher levels of pS129-α-Syn in ChAT neurons compared with wild type mice (Fig. 7, A, B).

**Fig. 7.**
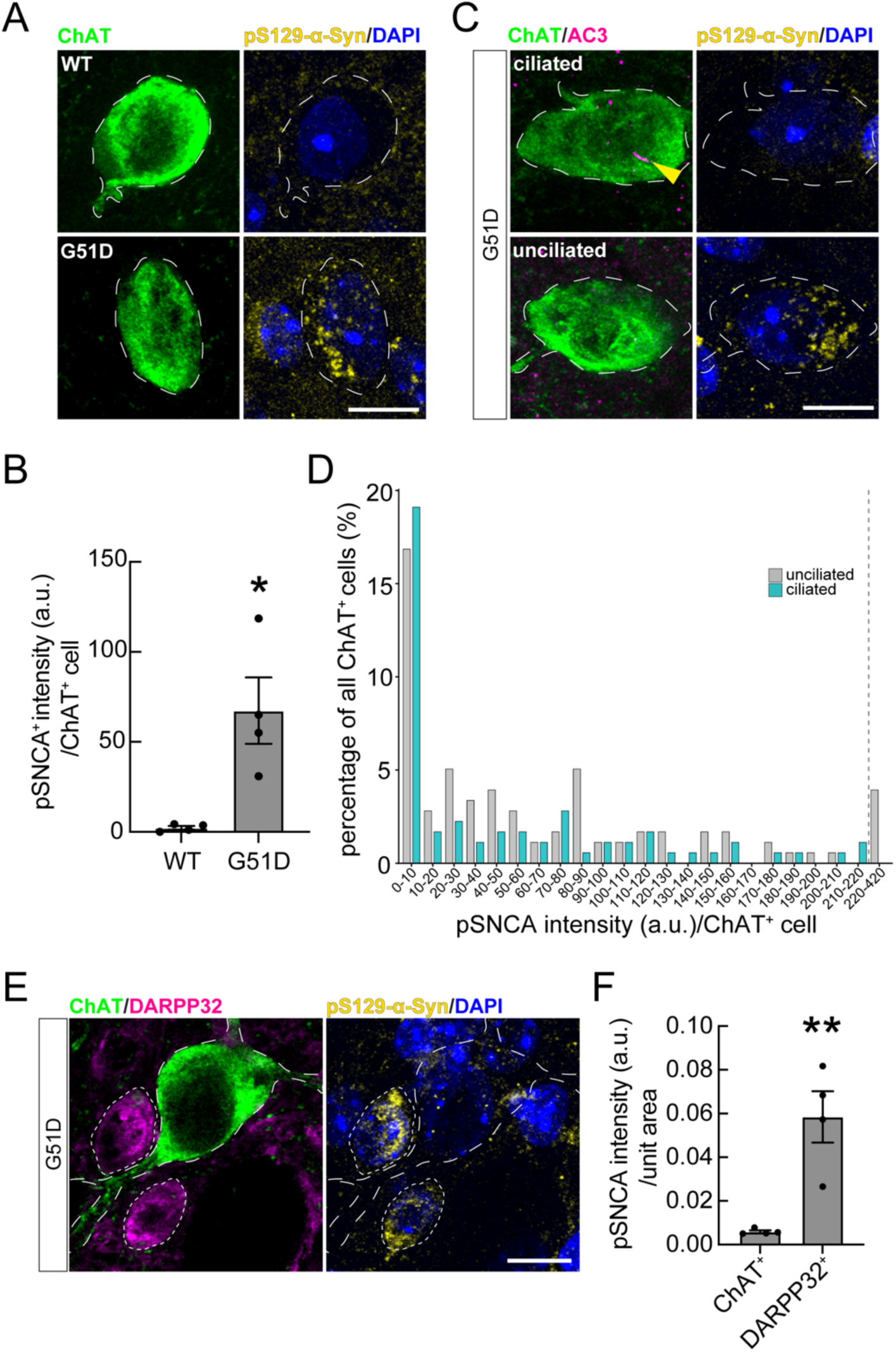
Loss of primary cilia correlates with increased phosphorylated α-synuclein (pSNCA) accumulation in striatal cholinergic neurons of *Snca^G51D/G51D^*mice. **(A)** Representative confocal immunofluorescence images of cholinergic neurons from 12-month-old wild type and *Snca^G51D/G51D^* mice. In all panels, cholinergic interneurons were labeled with anti-ChAT antibody (green), phosphorylated α-synuclein (pSNCA) was labeled with anti-pSNCA (Ser129) antibody (yellow), and nuclei with DAPI (blue). Scale bars, 10 μm. **(B)** Quantification of pSNCA⁺ intensity (arbitrary units) per ChAT⁺ interneuron. **(C)** Representative confocal images of ChAT neurons from 12-mo *Snca^G51D/G51D^*mice. Primary cilia were labeled with anti-AC3 antibody (magenta; yellow arrowheads). Scale bars, 10μm. **(D)** Single cell correlation between ciliation status and pSNCA intensity in ChAT⁺ neurons from*Snca^G51D/G51D^* mice. Single-cell pSNCA intensity was quantified per ChAT⁺ interneuron and binned, with cells plotted as the percentage of all ChAT⁺ neurons within each intensity range. Data are stratified by cilia status (ciliated, green; unciliated, gray). The dashed vertical line indicates a transition between fine and coarse binning. The analyzed population is the same as in Fig. 7B. **(E)** Representative confocal images of cholinergic and medium spiny neurons from 12-month *Snca^G51D/G51D^*mice. Cholinergic interneurons are circled (long dashed outlines); medium spiny neurons (MSNs) were labeled with anti-DARPP32 antibody (magenta, short-dashed outlines). Scale bars, 10 μm. **(F)** Quantification of the pSNCA⁺ intensity (arbitrary units, a.u.) per ChAT⁺ or DARPP32^+^ unit area from 12-month *Snca^G51D/G51D^* mouse brains. Values represent mean ± SEM from four mice per group (Fig. 7B) or per mouse per cell type (Fig. 7F). For Fig. 7B, two to three brain sections were analyzed per mouse, with >50 ChAT⁺ neurons scored per mouse. For Fig. 7F, neurons were quantified across 45 serial image sections and averaged to obtain one value per brain. Statistical significance was determined using an unpaired Student’s t test. *p = 0.0126; **p = 0.0043.

Quantification of pS129-α-Syn relative to ciliation status showed that, among striatal ChAT neurons, those with higher pS129-α-Syn levels were less likely to be ciliated (Fig. 7C, D). Panel 7D shows the distribution of pS129-α-Syn staining intensity across ChAT neurons in the G51D striatum and the corresponding ciliation status of cells within a given intensity bin. In G51D mouse striatum, approximately 45% of ChAT-positive neurons are ciliated, a value that is matched for ChAT neurons with 0-10 pS129-α-Syn staining units. Above this level of staining, the ration of unciliated to ciliated ChAT neurons increased.

In contrast, neighboring medium spiny neurons, which are far more abundant, retain their primary cilia in mouse and human PD models, including G51D (fig. S1), yet exhibit much higher pS129-α-Syn levels than ChAT neurons in the same striatal region (Fig. 7, E, F). Notably, at 12 months, the striatum overall has lower pS129–α-synuclein burden than other brain regions (*21*). Thus, pS129-α-Syn levels correlate with cilia loss within ChAT neurons but not within medium spiny neurons. This mirrors our previous finding that medium spiny neurons express more LRRK2 than ChAT neurons, yet ChAT neurons are more susceptible to cilia loss in mice expressing hyperactive, pathogenic LRRK2. Together, these data indicate that pS129-α-Syn levels do not predict overall vulnerability to cilia loss across different neuronal cell types.

## Discussion

G51D α-Syn mice provide a valuable model of idiopathic human PD, showing early olfactory deficits and age-dependent loss of motor function (*21*). We have shown here that G51D mice show cellular phenotypes shared by multiple genetic models of PD including mice and humans with pathogenic *LRRK2* mutations (*8, 12, 13, 15, 36*) or loss of the LRRK2-reversing PPM1H or PPM1M phosphatases (*9, 14*) and PINK1 knockout mice (*42*). These phenotypes include loss of primary cilia on cholinergic and parvalbumin neurons and ALDH1L1^+^ astrocytes (but not on the more abundant medium spiny neurons) in the dorsolateral striatum that are important for Hedgehog-dependent neurotrophic factor production in support of dopamine neurons. Although mouse models do not fully recapitulate human PD phenotypes such as significant loss of dopamine neurons in the substantia nigra and resting tremor, the G51D model has advantages in that it involves α-Syn expression from the endogenous mouse locus and provides spatiotemporal expression of mutant α-Syn that is analogous to PD.

G51D mice show initial deposition of α-Syn phosphorylation in the olfactory tract and vagal and enteric nerves, spreading along the olfactory circuit to the piriform cortex, entorhinal cortex, and hippocampus, and from the vagus nerve to the substantia nigra (*21*). The mice show delayed increases in pS129 α-Syn, which is not detectable at 3 months of age outside of the olfactory system and only becomes apparent at 12 months of age in deeper layers of the cortex, hippocampus, and substantia nigra (*21*). Mice display motor deficits from 9 months of age but do not show a significant reduction in dopamine levels until 18 months of age. These findings support the contention that early motor deficits are driven by dopaminergic neuronal dysfunction rather than absolute neuronal loss (*43*). In the striatum at 12 months of age, we detected increased pS129 α-Syn in G51D but not in wild type mice, with much more pronounced expression in medium spiny neurons than in ChAT neurons. Despite this distribution, ChAT neurons selectively lost primary cilia while the medium spiny neurons did not. Thus, important physiological changes in the striatum are unlikely to correlate directly with the overall pS129 α-Syn burden observed in this study.

Unlike what we have seen in LRRK2 mutant mice where GFAP^+^ astrocytes display a major loss of cilia as young as 10 weeks of age (*8*), G51D astrocytes appeared to require aging before a highly significant astrocytic ciliation difference was detected. Since astrocytes are filled with lysosomes, it is possible that this result can be explained by astrocytic capacity to degrade α-Syn aggregates (*44, 45*), explaining their relative decreased vulnerability to G51D α-Syn expression.

As discussed earlier, one of the first non-motor PD symptoms is loss of olfaction (*22, 46*). Odorants are first detected by peripheral olfactory sensory neurons of the olfactory epithelium. These neurons rely on their cilia for GPCR-mediated odorant detection (*29*) and given the impact of multiple PD-linked mutations on ciliary signaling, we were excited to explore OSN ciliary status in the G51D mice. We incorporated a protocol for isotropic expansion of the olfactory epithelium to enable super-resolution characterization of these multi-ciliated neurons. Under conditions of highly improved microscopic resolution, the cilia appeared fully intact on G51D olfactory sensory neurons. As mentioned earlier, the pathway to multi-ciliation is different from that used by cells containing a single primary cilium (*34, 35, 47*); nevertheless, it was important to evaluate olfactory sensory neuron ciliary status. In contrast, we did detect loss of primary cilia on horizontal basal cells of the olfactory epithelium that require cilia to enable them to regenerate damaged olfactory epithelia (*32*).

Upstream in the olfactory pathway, the piriform cortex integrates olfactory bulb inputs and forms widespread connections with other brain regions (*23, 24*). PV neurons within the piriform cortex regulate olfactory processing by sharpening sensory encoding and enhancing odor discrimination (*24, 25*), and their excitability is increased by dopamine (*48*). Functional MRI studies show decreased activation and weaker integration with the broader olfactory–network during odor stimulation in PD compared with healthy controls (*49*), implicating the piriform circuits as a key locus of PD-related olfactory disruption. We observed ciliary loss and reduced NRTN production in piriform PV neurons, suggesting that cilia loss or dysfunction may impair PV neuron activity, local interneuron maintenance or survival. Future work will reveal the full consequences of piriform PV neuron cilia loss and corresponding decrease in NRTN production.

Idiopathic PD is thought to be driven by α-Syn pathology (*50*). We previously reported cell-type specific cilia loss in postmortem striatal tissue from patients with idiopathic PD (*12, 13*), consistent with the proposal that cell type-specific ciliation is influenced by α-Syn pathology. Our findings with G51D mice support this conclusion. How might mutant α-Syn trigger cilia loss? One possible explanation would be that α-Syn aggregates activate LRRK2, a kinase that regulates ciliogenesis and is activated by lysosome stress (*38, 51*). It will be important to evaluate the phenotypes of LRRK2^-/-^ or LRRK2-inhibited G51D α-Syn mice, to test this hypothesis. Prior work showed that hyperactive LRRK2 or LRRK2 overexpression exacerbated synuclein pathology (*52, 53*) but striatal and piriform cortex ciliation was not evaluated. Independent of LRRK2, ciliation is linked to induction of autophagy (*54, 55*); α-Syn aggregates might overload the autophagy machinery (*56*) and in that manner, somehow disrupt ciliogenesis. Future work will be needed to explain the molecular basis of LRRK2-dependent or independent ciliogenesis blockades and the resistance of medium spiny neurons to cilia loss in both mouse and human.

In addition to controlling primary ciliogenesis, LRRK2 catalyzes lysosome exocytosis, especially in cells of the macrophage lineage (*57–59*). While exocytic release of lysosomal α-Syn aggregates may be beneficial for neurons, microglial uptake of these aggregates and their subsequent (LRRK2-stimulated) exocytosis may also contribute to unfavorable α-Syn spread. Thus, LRRK2 may protect neurons in the short term but contribute to disease pathology in the longer term. Future experiments with the G51D α-Syn mouse model should be able to provide important information related to whether LRRK2 inhibition is beneficial or instead, exacerbates α-Syn pathology in the brain. Generation of mice with cell type-specific expression of LRRK2 may also help determine the cells in which it plays the most significant role, be it a subset of neurons, astrocytes, microglia or all three.

Hedgehog signaling is critical for dopamine neuron survival (*17*) and 3 months of LRRK2 inhibitor feeding can restore primary cilia, Hedgehog signaling and downstream, striatal neuroprotective factor production in *Lrrk2* mutant mice (*15*). These findings hold great promise for patients with LRRK2 mutations and encourage therapeutic development of this class of small molecule LRRK2 inhibitors. It will be very important to understand the contribution of LRRK2 hyperactivity to idiopathic PD pathophysiology to determine whether LRRK2 inhibitors will be beneficial to the larger number of patients with idiopathic PD.

In summary, we have found mice expressing G51D α-Syn share vulnerability to cilia loss in the striatum with mice carrying pathogenic LRRK2 mutations or with PPM1H, PPM1M or PINK1 gene knockouts: striatal cholinergic and parvalbumin interneurons, as well as astrocytes, lose primary cilia and the neurotrophic signaling needed to sustain dopaminergic neurons. We showed that these vulnerable cells do not contain higher levels of pS129 α-Syn than the surrounding medium spiny neurons that contain significantly more. We showed previously that striatal medium spiny neurons express more LRRK2 RNA than the vulnerable neurons, but in a given class of neuron, those with higher LRRK2 showed increased probability of cilia loss (*12*). Despite sharing a similar phenotype, the timing of cilia loss differed among these models. For example, astrocyte cilia loss was only truly notable at 12 months in G51D mice while it is readily observed in LRRK2 pathway mutant mice at 10 weeks of age. This suggests that LRRK2 is a more direct regulator of ciliogenesis than α-Syn aggregates. Yet the ages of onset for LRRK2 carriers are not so different from that of patients with idiopathic disease. Perhaps this is because apparently earlier-acting LRRK2 mutations impact about half of cholinergic and parvalbumin interneurons at all ages, while later acting α-Syn may more slowly, eventually, impact all. Altogether, these experiments highlight the importance of Hedgehog signaling in multiple brain regions for neuroprotection, especially for dopamine neuron survival. And cilia must also play other fundamental roles for neuronal regulation, given their abilities to synapse directly upon one another (*60, 61*).

## Materials and Methods

### Research standards for animal studies

Mice were housed in a level 3, American Association for Laboratory Animal Science–certified facility on a 14-h light cycle. Husbandry, housing, euthanasia of animals, and experimental guidelines were approved by the Institutional Animal Care & Use Committee (IACUC) at Baylor College of Medicine.

The *Snca^G51D/G51D^*knock-in mice were generated at Baylor College of Medicine; genotyping was confirmed by genomic polymerase chain reaction with different primers between the unmodified wild type and modified G51D alleles (*21*).

### Mouse brain and olfactory epithelium processing

*Snca^G51D/G51D^* mouse brains (3- and 12-month-old) and age-matched wild type controls were fixed by transcardial perfusion using 4% paraformaldehyde (PFA) in PBS as described dx.doi.org/10.17504/protocols.io.bnwimfce. Following perfusion, whole brains were extracted, and the olfactory epithelium was rapidly dissected as described (*62*). Both brains and olfactory epithelium were post-fixed in 4% PFA for 24 hr and then immersed in 30% (w/v) sucrose in PBS until the tissue settled to the bottom of the tube (∼48 hr). The brains and olfactory epithelium were harvested at Baylor College of Medicine and sent to Stanford University for analysis. Prior to cryosectioning, tissues were embedded in cubed-shaped plastic blocks with OCT (BioTek, USA) and stored at −80°C. OCT blocks were allowed to reach −20°C for ease of sectioning. The brains and olfactory epithelium were oriented to cut coronal sections on a cryotome (Leica CM3050S, Germany) at 20 μm thickness for brain tissue and 14 µm thickness for olfactory epithelium and positioned onto SuperFrost plus tissue slides (Thermo Fisher, USA).

### Mouse tissue homogenization and immunoblotting analysis

*Snca^G51D/G51D^* (9-months-old) and age-matched wild type controls were euthanized, and whole brains were harvested and immediately frozen on dry ice. Frozen brain tissues were collected at Baylor and shipped on dry ice to Stanford for downstream analysis. Frozen brain tissues were homogenized as described in dx.doi.org/10.17504/protocols.io.sqgedtw. Briefly, frozen brain tissues were homogenized using M tubes in a GentleMACS machine with lysis buffer (50 mM Tris–HCl pH 7.4, 150 nM NaCl, 1 mM EGTA, 2% glycerol, cOmplete EDTA-free protease inhibitor cocktail (Roche), PhosSTOP phosphatase inhibitor cocktail (Roche), 1 μg/ml microcystin-LR (Sigma), and 1% (v/v) Triton X-100). 6 mL of lysis buffer was used per brain, about 500-600 mg frozen weight each. Tissues were homogenized using the protein cycle, run twice with 5 mins incubation on ice between each cycle.

Following homogenization, M-tubes were spun briefly at 2000 x g at 4°C for 5 minutes to pellet any unhomogenized material. The supernatant was divided into 1.5 mL Eppendorf tubes, then further clarified by centrifugation at 20,000 x g at 4°C for 30 minutes.

Lysate protein was determined by Bradford assay. Samples containing equal protein amounts were mixed with 5x SDS sample buffer (250 mM Tris-HCl, pH 6.8, 30% glycerol (v/v), 10% SDS (w/v), 0.1% bromophenol blue (w/v), 10% 2-mercaptoethanol (BME) (v/v)) and resolved on 4-20% precast gels (Bio-Rad) then transferred onto nitrocellulose membranes using the Transblot Turbo System (Bio-Rad). Membranes were blocked in 3% BSA in Tris-buffered saline (200 mM Tris, 1.5M NaCl) with 0.1% Tween-20 (TBST) for 30 minutes and incubated with specific primary antibodies overnight at 4°C.

See Key Resource Table (table S1) for primary and secondary antibodies used and their dilutions. Primary antibodies were diluted in 3% BSA in TBST and detected using LI-COR IRdye labeled secondary antibodies (donkey anti-mouse 680/800, donkey anti-rabbit 680/800), diluted 1:10,000 in 3% BSA in TBST. Membranes were scanned on a LICOR Odyssey DLx scanner. Images were saved as .tif files and analyzed using the gel scanning plugin in FIJI (*63*).

### Immunohistochemical staining

Mouse brain striatum and olfactory epithelium were subjected to immunostaining following a previously established protocol (dx.doi.org/10.17504/protocols.io.bnwimfce). Frozen slides were thawed at room temperature for 15 minutes and then gently washed twice with PBS for 5 minutes each. Antigen retrieval was achieved by incubating the slides in 10 mM sodium citrate buffer pH 6.0, preheated to 95°C, for 15 minutes. Sections were permeabilized with 0.1% Triton X-100 in PBS at room temperature for 15 minutes, followed by blocking with PBS containing 2% FBS and 1% BSA for 2 hr at room temperature. Primary antibodies were applied overnight at 4°C, and sections were then exposed to secondary antibodies at room temperature for 2 hr. Secondary antibodies used were donkey highly cross-absorbed H + L antibodies conjugated to Alexa 488, Alexa 568 and Alexa 647 diluted at 1:2000. Nuclei were counterstained with 0.1 μg/ml DAPI (Sigma). Finally, stained tissues were mounted with Fluoromount G and covered with a glass coverslip. All antibody dilutions for tissue staining contained 1% DMSO to facilitate antibody penetration.

### Expansion microscopy of olfactory epithelium

Expansion microscopy of the olfactory epithelium was performed as described (dx.doi.org/10.17504/protocols.io.6qpvry22ogmk/v1) to achieve ∼4-5X isotropic expansion. Frozen olfactory epithelium sections were thawed at room temperature for 1 minute, washed three times in PBS, and incubated for 2 hr at RT in a monomer-fixative solution containing 0.7% PFA and 1% acrylamide in PBS to anchor proteins to the gel matrix. Sections were then embedded in a hydrogel composed of 19% sodium acrylate, 10% acrylamide and 0.1% bis-acrylamide in PBS. Polymerization was initiated using ammonium persulfate and TEMED, first at 4°C for 5 minutes and subsequently overnight at room temperature in a humidified chamber. Gels were denatured at 95°C for 6 hr in a buffer containing 50 mM Tris (pH 9.0), 200 mM NaCl, and 200 mM SDS. Gels were then fully expanded by repeated incubation in deionized water. Expanded gels were incubated overnight at 4°C with goat anti-OMP (1:100) and mouse anti-acetylated tubulin (1:500), followed by overnight incubation at 4°C with Alexa Fluor conjugated secondary antibodies (1:500) and DAPI. The gels were washed in PBS, fully expanded in deionized water and mounted on poly-D-lysine coated imaging dishes and imaged using Zeiss LSM 900 confocal microscope to visualize olfactory sensory neuron cilia.

### Fluorescence in situ hybridization (FISH)

RNAscope fluorescence in situ hybridization was carried out as described (https://bio-protocol.org/exchange/preprintdetail?id=1423&type=3&searchid=EM1708992000021453&sort=5&pos=b; (Khan et al., 2021)). The RNAscope Multiplex Fluorescent Detection Kit v2 (Advanced Cell Diagnostics) was utilized following the manufacturer’s instructions, employing RNAscope 3-plex Negative Control Probe (#320871) or Mm-Gdnf (#421951), Mm-Nrtn-C2 (#441501-C2), Mm-Pvalb-C3 (#421931-C3) and Mm-Bdnf-C1 (#424821). The Mm-GDNF, Mm-Nrtn-C2, and Mm-Pvalb-C3 probes were diluted 1:30, 1:4 and 1:10, respectively in dilution buffer consisting of 6x saline-sodium citrate buffer (SSC), 0.2% lithium dodecylsulfate, and 20% Calbiochem OmniPur Formamide. Fluorescent visualization of hybridized probes was achieved using Opal 690 or Opal 570 (Akoya Biosciences).

Subsequently, brain slices were subjected to blocking with 1% BSA and 2% FBS in TBS (Tris buffered saline) with 0.1% Triton X-100 for 30 minutes. They were then exposed to primary antibodies overnight at 4°C in TBS supplemented with 1% BSA and 1% DMSO. Secondary antibody treatment followed, diluted in TBS with 1% BSA and 1% DMSO containing 0.1 μg/ml DAPI (Sigma) for 2 hr at room temperature. Finally, sections were mounted with Fluoromount G and covered with glass coverslips.

### Microscope image acquisition

All images were obtained using a Zeiss LSM 900 confocal microscope (Axio Observer Z1/7) coupled with an Axiocam 705 camera and immersion objective (Plan-Apochromat 63x/1.4 Oil DIC M27). The images for epithelial cells, astrocytes and neurons were acquired at 0.4, 0.5, and 0.75 µm Z-sampling, respectively. The images were acquired using ZEN 3.4 (blue edition) software, and visualizations and analyses were performed using ImageJ Fiji (*63*), CellProfiler (*64*) and R (65).

### Quantification of phospho-α-Synuclein in mouse striatal cholinergic neurons and analysis by cilia status

Fluorescence microscopy z-stacks from wild type and*Snca*^G51D/G51D^ mutant mouse brain tissues stained for pSNCA, AC3, ChAT and DAPI were acquired and processed in FIJI/ImageJ using a custom FIJI macro (dx.doi.org/10.17504/protocols.io.5jyl8xjzdv2w/v1) to generate non-overlapping 3-slice maximum-intensity projections across the full z-depth for each channel. All 3-slice projections derived from the same z-stack were treated as belonging to a single image dataset corresponding to one field of view. Projection images were imported into CellProfiler (*64*), where pSNCA-positive signal was segmented and restricted to only ChAT-positive cholinergic neurons. Integrated pSNCA intensity was measured for each ChAT-positive cholinergic neuron. A detailed protocol of the CellProfiler (*64*) pipeline is referenced here (dx.doi.org/10.17504/protocols.io.261ge13bwv47/v2).

For comparisons within the *Snca*^G51D/G51D^ mutant group, pSNCA measurements were grouped according to cilia status. Ciliary status (present or absent) was annotated (dx.doi.org/10.17504/protocols.io.bnwimfce) for each ChAT-positive neuron and linked to the corresponding pSNCA measurement from the same segmented cell. CellProfiler output tables were imported into R (65) for analysis. pSNCA distributions were visualised using histograms, plotted as the percentage of neurons per intensity bin.

### Quantification of phospho-α-Synuclein in mouse striatal cholinergic and medium spiny neurons

Fluorescence microscopy z-stacks from *Snca*^G51D/G51D^ mutant mouse brain tissues stained for pSNCA, DARPP32, ChAT, and DAPI channels, were processed using the same custom FIJI/ImageJ macro described above (dx.doi.org/10.17504/protocols.io.5jyl8xjzdv2w/v1) to generate non-overlapping 3-slice maximum-intensity projections across the z-axis. Projection images were imported into CellProfiler, where pSNCA-positive signal was segmented and cell-type masks were generated using DARPP32 and ChAT fluorescence to identify medium spiny neurons (MSNs) and ChAT-positive cholinergic neurons, respectively. Integrated pSNCA intensity was measured within each cell-type mask for every 3-slice projection. The same protocol described above (dx.doi.org/10.17504/protocols.io.261ge13bwv47/v2) was used for analysis, in which integrated pSNCA fluorescence intensities were summed per dataset separately for ChAT-positive and DARPP32-positive neurons and normalised to the corresponding total segmented cell mask area, yielding cell type–specific pSNCA intensity per unit area.

All scripts (macro and R) and the CellProfiler pipeline are available at https://doi.org/10.5061/dryad.4xgxd25r0.

## Statistical analysis

In addition to above-mentioned methods, all other statistical analysis was carried out using GraphPad Prism version 10.2.3 for Macintosh, GraphPad Software, Boston, MA, USA, www.graphpad.com.

## Acknowledgments

Funding: This research was funded by Aligning Science Across Parkinson’s (ASAP-000463 to SRP) through the Michael J. Fox Foundation for Parkinson’s Research (MJFF) and HHMI and Freedom Together Foundation (to HYZ). For the purpose of open access, the author has applied a CC BY public copyright license to all Author Accepted Manuscripts arising from this submission.

## Author contributions

**Conceptualization: Y-E.L., E.J., H.Y.Z., S.R.P.**

**Mice, Tissue dissections: Y.D.K., B.D.W.B., E.J., H.Y.Z.**

**Methodology: Y-E.L., E.J, A.V., B.D.W.B.**

**Investigation: Y-E.L., E.J., C.Y.C., A.L.**

**Visualization: Y-E.L., E.J., S.R.P., C.Y.C., A.V.**

**Supervision: S.R.P., H.Y.Z., B.R.A.**

**Writing—original draft: S.R.P. and Y-E.L.**

**Writing—review & editing: All authors**

## Data and materials availability

The data, code, protocols, and key lab materials used and generated in this study are listed in a Key Resources Table alongside their persistent identifiers at https://doi.org/10.5061/dryad.4xgxd25r0. All data needed to evaluate the conclusions in the paper are present in the paper or the Supplementary Materials.

---

**Fig. S1.**
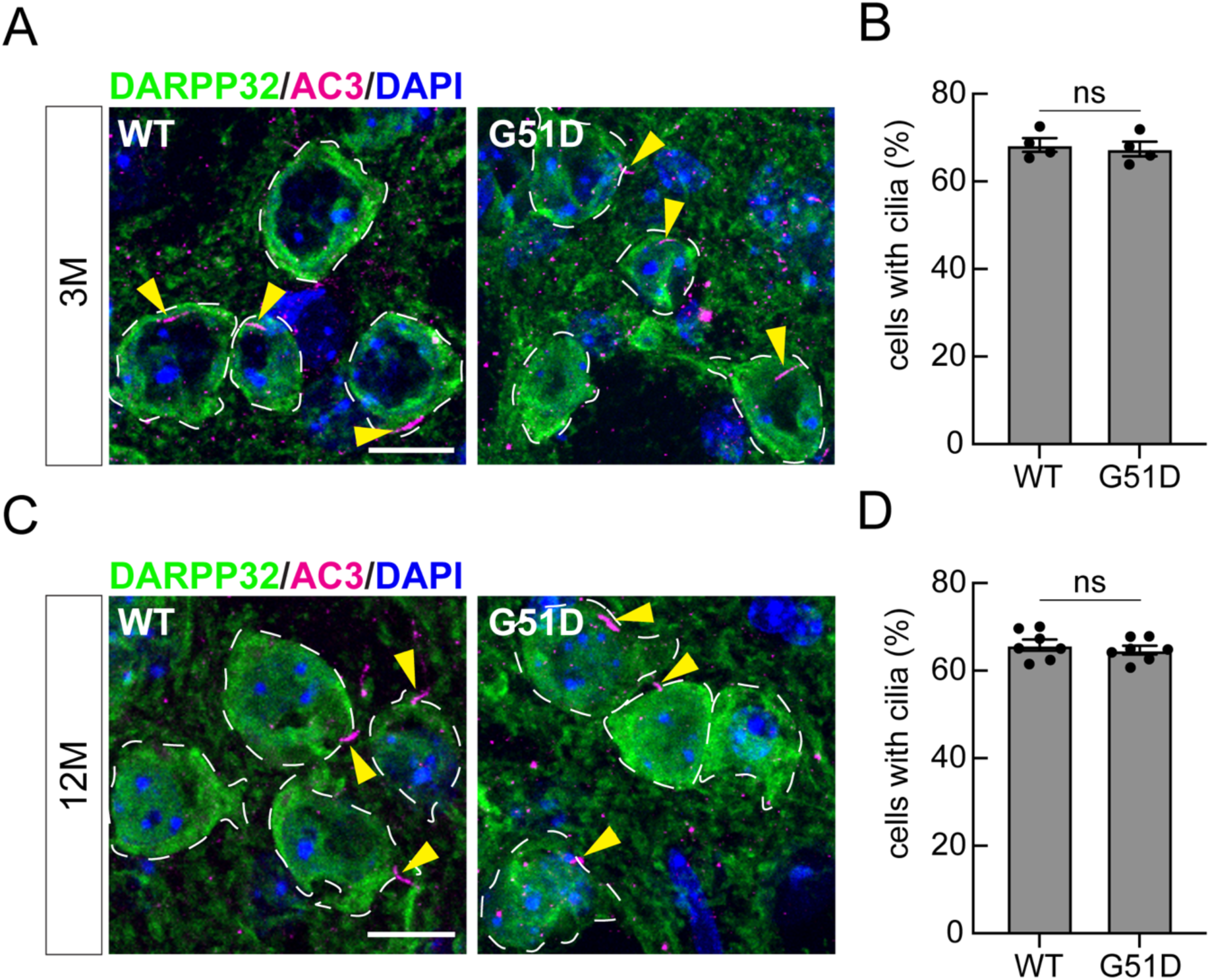
The *Snca^G51D/G51D^*mutation does not induce cilia defects in predominant medium spiny neurons of the mouse dorsal striatum. **(A and C)** Representative confocal immunofluorescence images of dorsal striatal sections from 3- and 12-month-old wild type and *Snca^G51D/G51D^* mice. Medium spiny neurons (MSNs) were labeled with anti-DARPP32 antibody (green). Primary cilia were labeled with anti-AC3 antibody (magenta; yellow arrowheads). Nuclei were labeled with DAPI (blue). Scale bars, 10 μm. **(B and D)** Quantitation of DARPP32^+^ neurons containing a cilium in 3- and 12-month-old wild type and *Snca^G51D/G51D^*mice, respectively. Values represent mean ± SEM from four (3-month-old) or seven (12-month-old) individual mice per group, with two to three sections analyzed per mouse. More than 300 DARPP32^+^ MSNs were scored per mouse. Statistical significance was determined using an unpaired Student’s *t* test. Differences were not significant (ns; p = 0.7093 at 3 months and p = 0.4837 at 12 months).

**Fig. S2.**
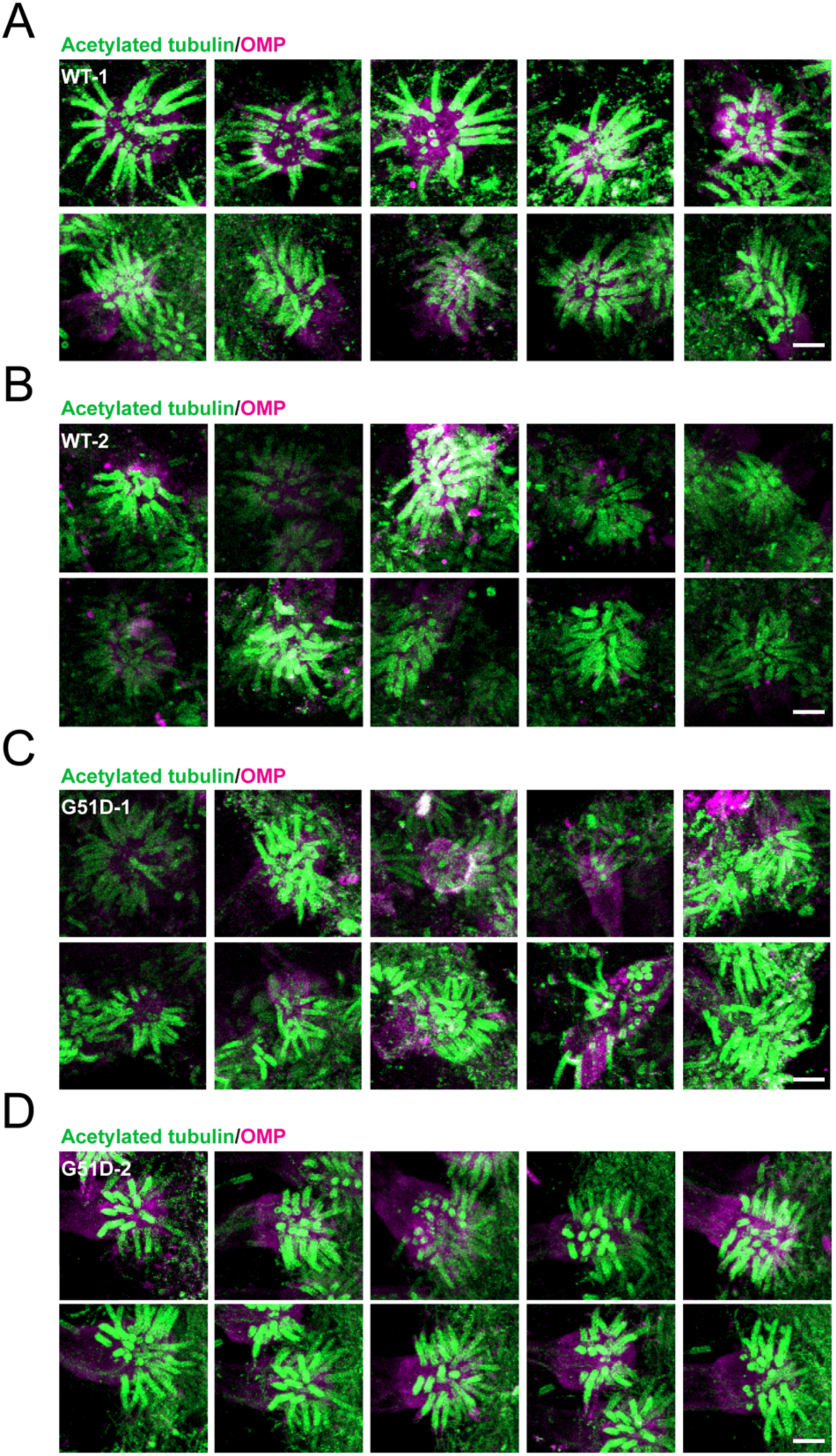
Representative confocal images of the OSN cilia of 12 month-old wild type and *Snca^G51D/G51D^* mice visualized using expansion microscopy. OSNs were labeled with anti-OMP antibody (magenta), and cilia with anti-Acetylated alpha-tubulin antibody (green). All images are from 12 month-old animals. (A) Wild type animal 1(WT-1), (B) Wild type animal 2 (WT-2), (C) *Snca^G51D/G51D^*animal 1 (G51D-1). (D) *Snca^G51D/G51D^* animal 2 (G51D-2). Scale bar, 1 µm.

**Fig. S3.**
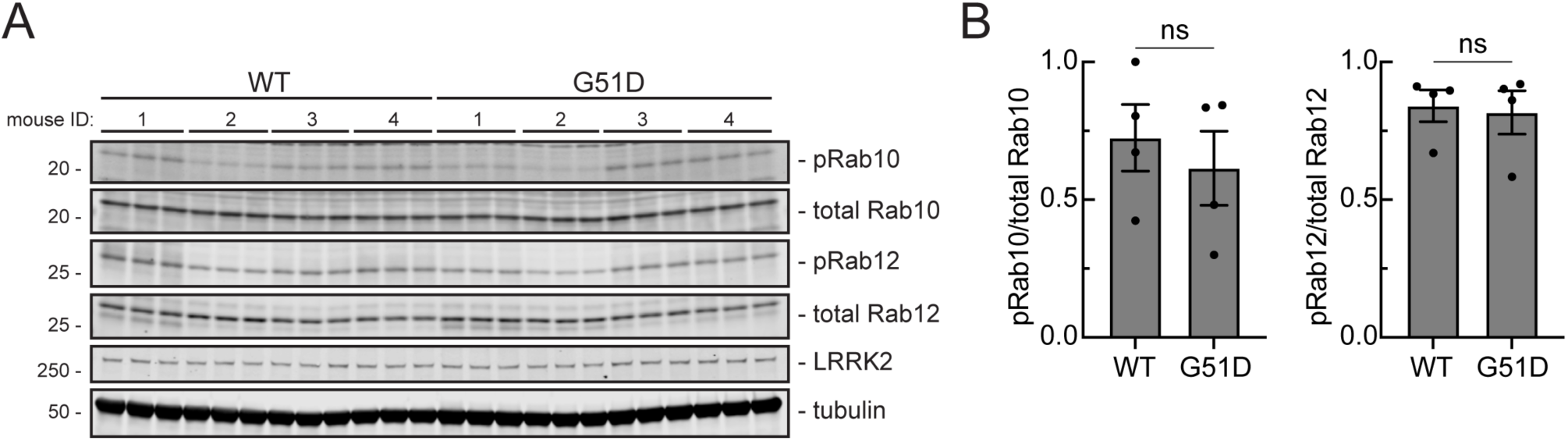
The *Snca^G51D/G51D^*mutation does not affect phosphorylated Rab levels in whole brain tissue. **(A)** Immunoblot analysis of 9 month-old wild type (WT) and *Snca^G51D/G51D^* mouse whole brain lysate. Four mice were analyzed per genotype (mouse ID 1-4). Mass is shown on the left in kDa; antigens are indicated at the right. **(B)** Quantitation of phosphoRab10 (pRab10)/total Rab10 and phosphoRab12 (pRab12)/total Rab12 levels from immunoblots in (A). Values were normalized to 1.0 for the highest WT level, and each dot represents the average of two independent replicates from one mouse.

**Table S1.**
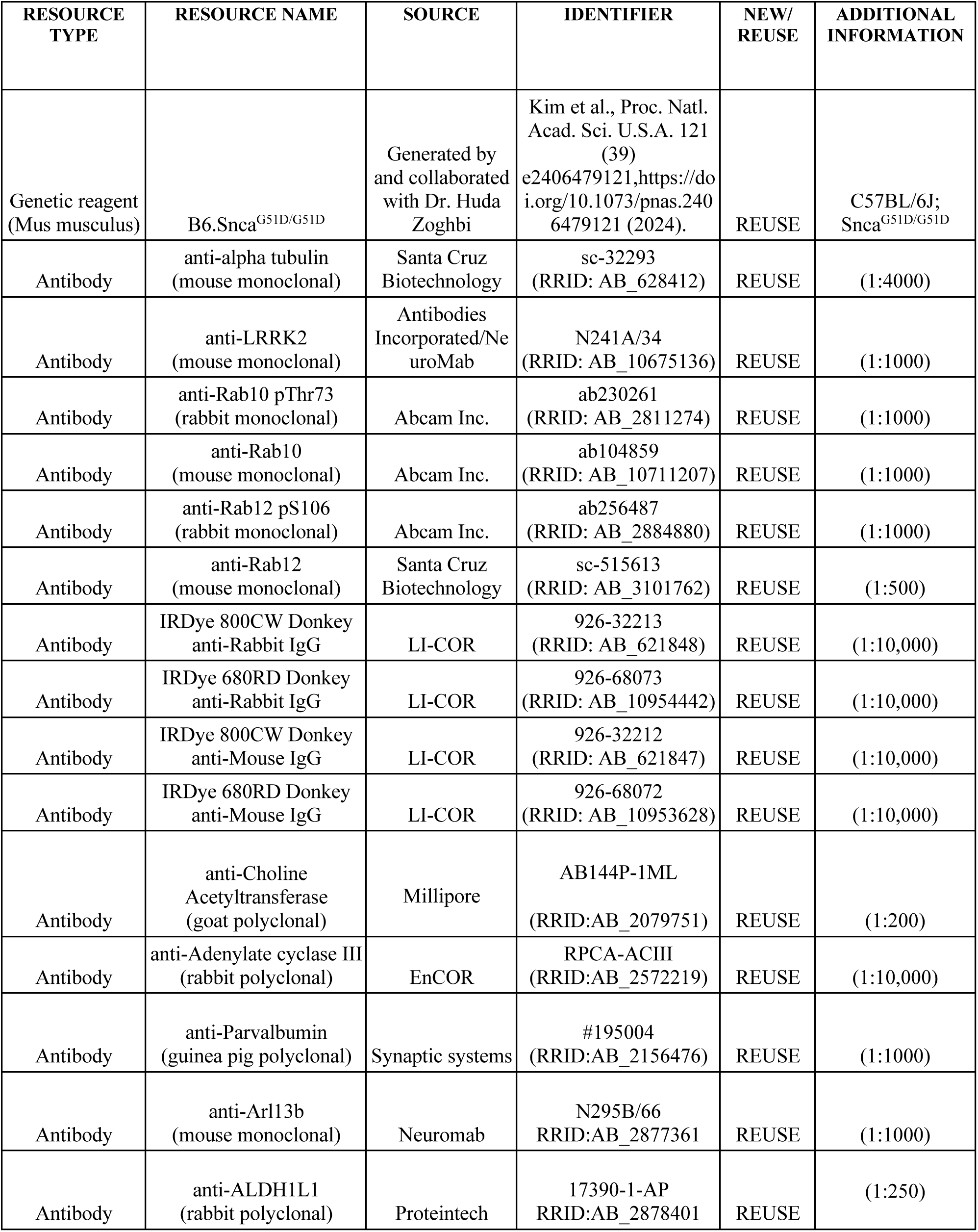
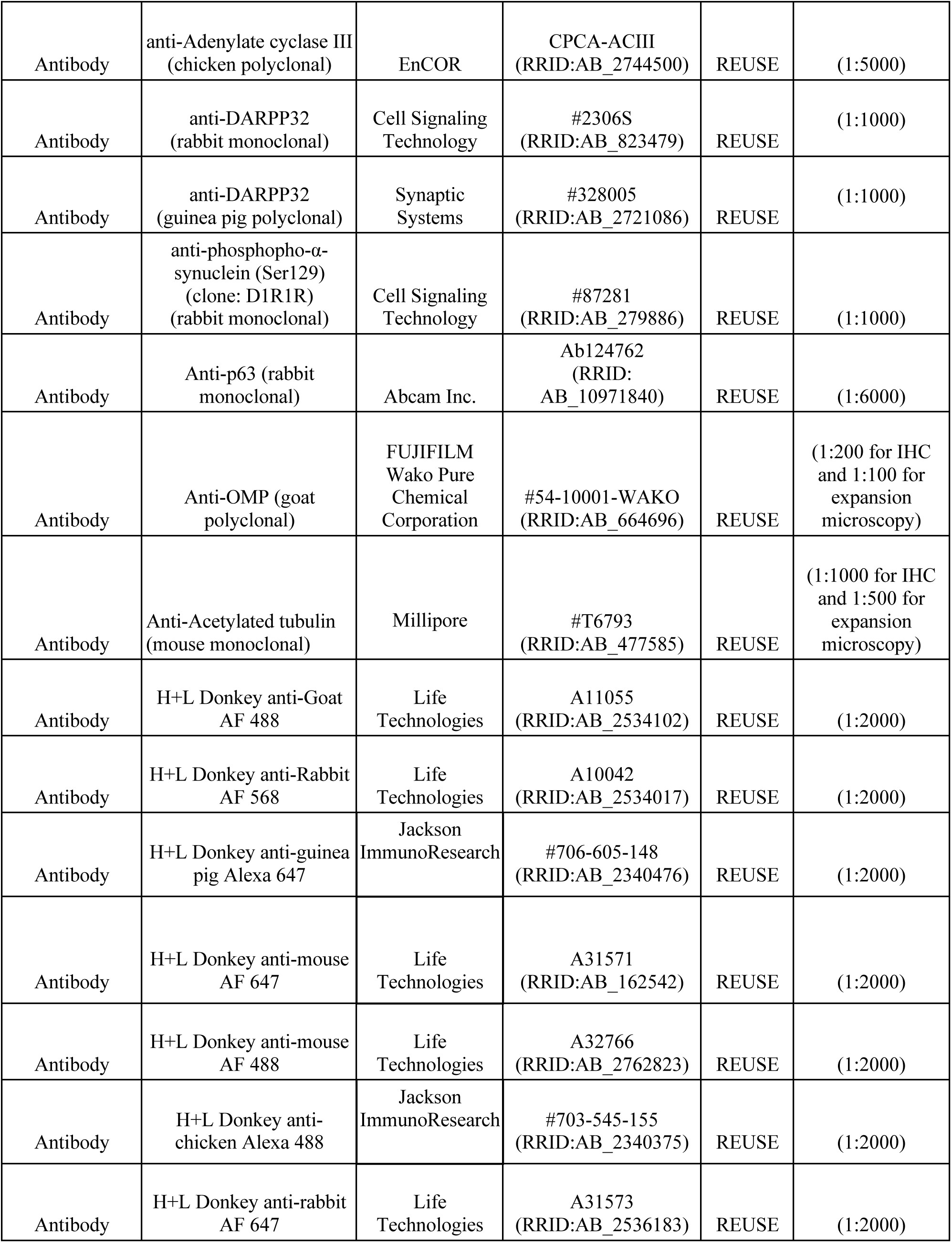
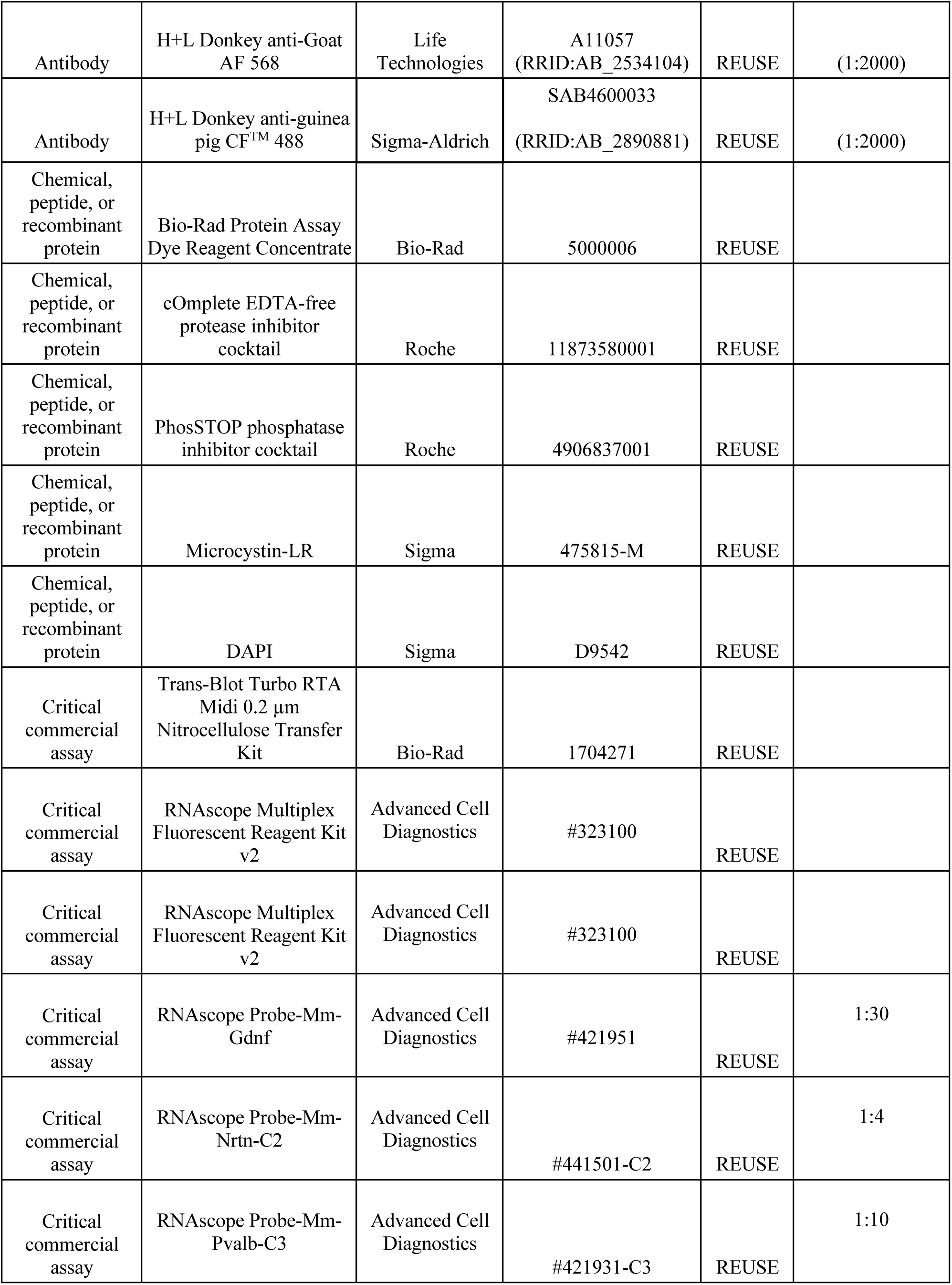
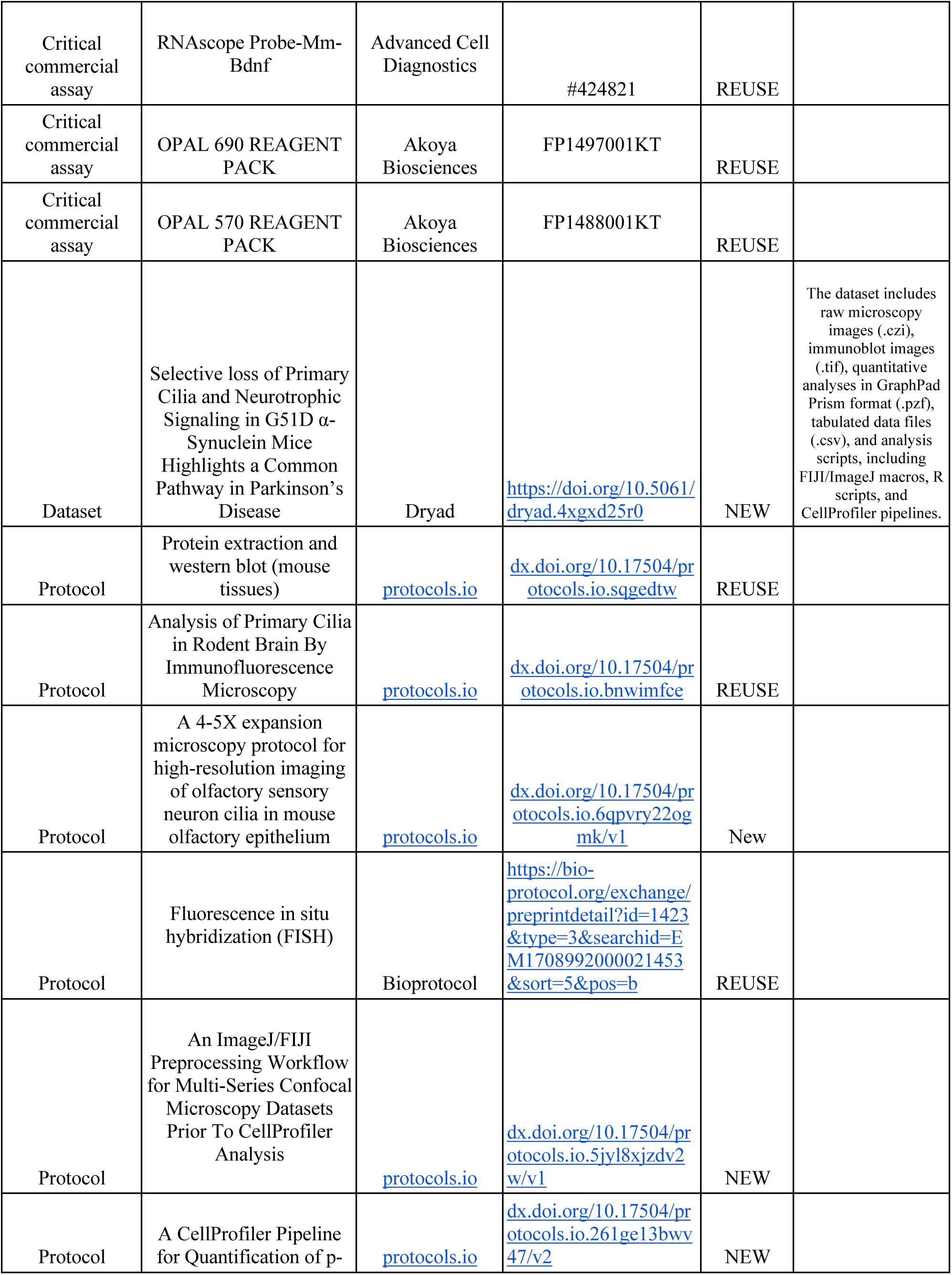
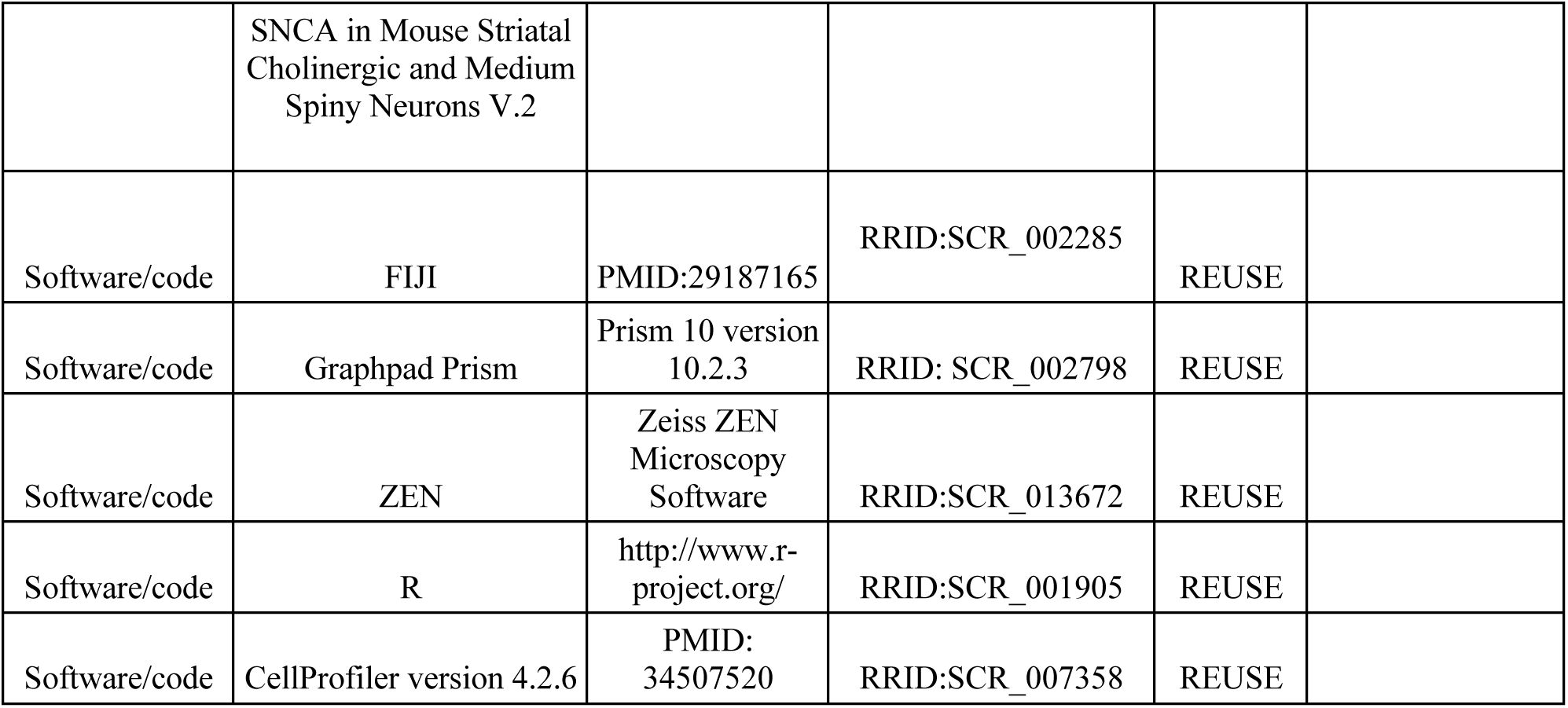
Key Resource Table.

